# Hipk is required for JAK/STAT activity and promotes hemocyte-derived tumorigenesis

**DOI:** 10.1101/058156

**Authors:** Jessica A. Blaquiere, Nathan B. Wray, Esther M. Verheyen

## Abstract

Dysregulation of key signaling molecules and pathways are causative of many Human diseases and cancers. A point mutation in the Drosophila Janus kinase (called *hop*) causes constitutive activation of the JAK/STAT pathway and results in blood cell tumours. We provide robust genetic evidence that Hipk is required for endogenous JAK/STAT activity. Overexpression of Hipk can phenocopy the effects of overactive JAK/STAT mutations and lead to melanized tumors and loss of Hipk can suppress the effects of hyperactive JAK/STAT. Furthermore, Hipk expression in blood cell progenitors causes tumors. PLA experiments show that Hipk can interact with the pathway effector Stat92E. Together our results show that Hipk is a novel factor required for effective JAK/STAT signaling.

**Summary Statement:** Loss of *hipk* impairs JAK/STAT activity in multiple tissue types and elevated Hipk leads to the formation of blood cell tumors in Drosophila.

## Introduction

Drosophila is a useful model to study evolutionarily conserved signaling pathways that are used reiteratively during development, as well as for modeling diseases, such as leukemia. Dysregulation of the JAK/STAT pathway has been linked to leukemia, myeloproliferative neoplasms, and solid tumors in flies and vertebrates (Amoyel et al., 2014; Dearolf, 1998; Jones et al., 2005; Lacronique, 1997; Levine et al., 2005). In Drosophila, the core components of the pathway include the Unpaired ligands (Upd, Upd2, Upd3), the Domeless receptor (Dome), the JAK homolog Hopscotch (Hop), and the transcription factor Stat92E (reviewed in Chen et al., 2014). Upon cascade stimulation, Stat92E becomes phosphorylated, dimerizes, and travels to the nucleus to regulate JAK/STAT target genes. JAK/STAT mutations are heavily correlated with tumor invasiveness and lethality (Hanratty and Dearolf, 1993). *hop^Tum-I^* is a dominant mutation resulting in a hyperactive Hop kinase that leads to constitutive activation of the pathway (Harrison et al., 1995). Similar activating JAK2 mutations are commonly seen in vertebrate cancers (Jones et al., 2005; Kralovics et al., 2005).

Homeodomain-interacting protein kinase (Hipk in Drosophila, Hipk1-4 in vertebrates) regulates numerous conserved signaling pathways (Chen and Verheyen, 2012; Lee et al., 2009a; Lee et al., 2009b; Poon et al., 2012; Rinaldo et al., 2008; Swarup and Verheyen, 2011). Interestingly, Hipk overexpression results in tumor-like formations similar to those seen in *hop^Tum-l^* flies, prompting our investigation into Hipk’s role in the JAK/STAT pathway. Indeed, reducing *hipk* suppressed the severity of the *hop^Tum-l^* phenotype. Further, we provide evidence that Hipk is required for JAK/STAT activity in a kinase-dependant manner and that Hipk and Stat92E interact *in vivo*. Our data indicate a novel role for Hipk in regulating JAK/STAT activity in endogenous and tumorous conditions.

## Results and Discussion

### Hipk induces hemocyte-derived melanotic tumors

We observed that Hipk induces pigmented masses, a phenotype resembling flies with overactive JAK/STAT signaling (Hanratty and Ryerse, 1981; Luo et al., 1995). Overexpression of *hipk* with *dpp-GAL4* (*dpp>HA-hipk^3M^*+2x*GFP*) caused the formation of pigmented tumors in the larval stages of development (Fig. 1B). These masses were not due to cell death, since they persisted when cell death was blocked with P35 (Fig. 1C) (Hay et al., 1994). Melanotic tumors induced by *hop^Tum-l^* arise due to overamplification and melanization of hemocytes, fly hematopoietic cells (Hanratty and Ryerse, 1981). Therefore, we next tested whether *hipk* could cause tumors when overexpressed in the circulating hemocytes and lymph gland using *hemolectin-GAL4* (*hml-GAL4*) (Sinenko and Mathey-Prevot, 2004). 91.7% of *hml>HA-hipk^3M^* flies exhibited at least one melanotic tumor, with the average being 3-4 tumors (Fig. 1E,F), compared to 0% of *hml>GFP* flies (Fig. 1D).

**Figure 1.**
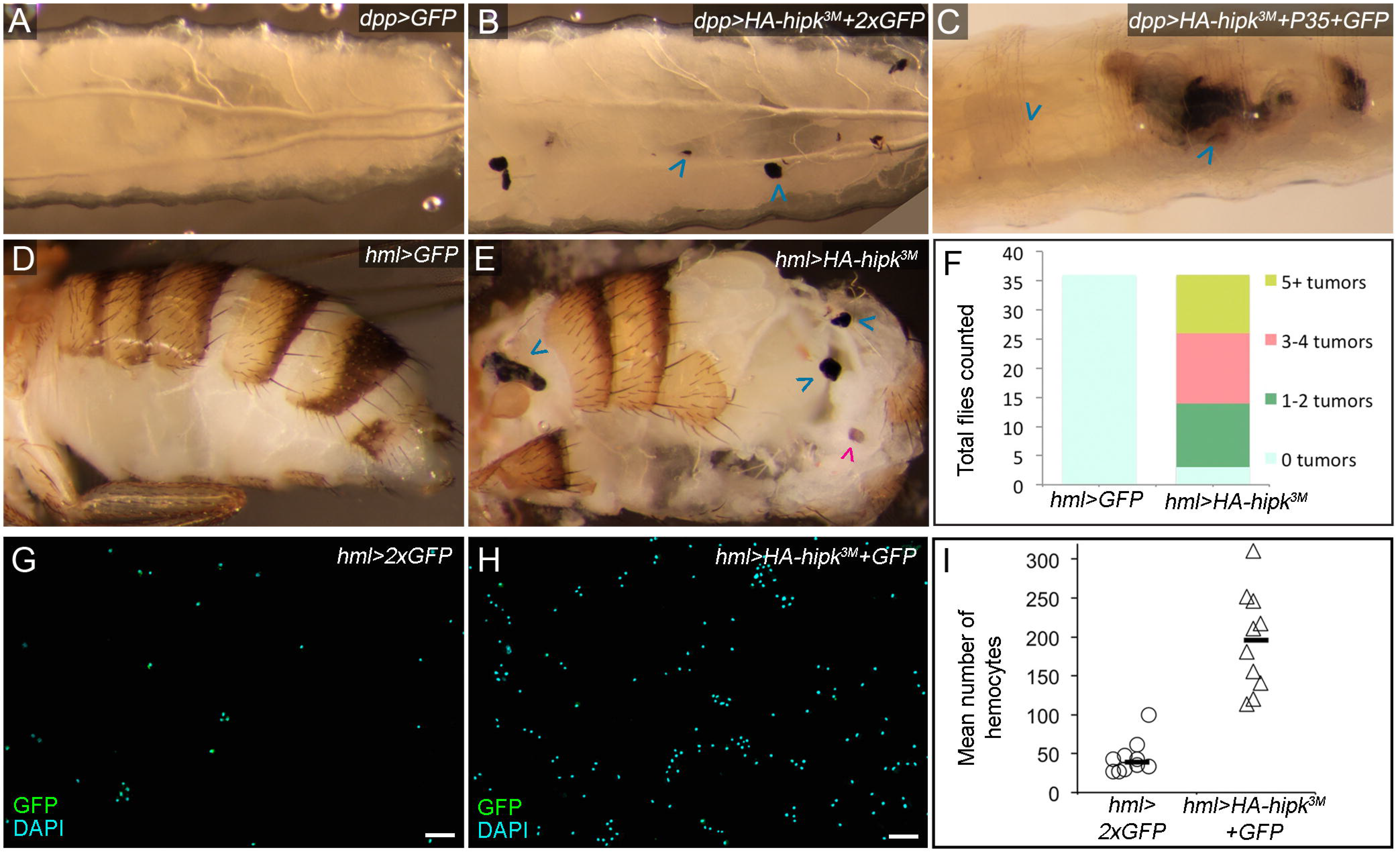
Hipk induces hemocyte derived melanotic tumors. (**A**) A control *dpp>GFP* L3 larva. (**B**) Stationary melanized masses are observed in 65% of *dpp>HA-hipk*^3M^+*GFP* larvae (blue arrowheads; n=40) and (**C**) persist when apoptotic cell death is inhibited in *dpp>HA-hipk^3M^+P35+GFP* larvae. (**D**) The abdomen of a control *hml>GFP* fly. (**E**) Melanized tumors are present in *hml>HA-hipk^3M^* flies (blue arrowheads). Spermathecae were not counted (magenta arrowhead). (**F**) Quantification of the number of tumors scored in from the dissected abdomens of flies shown in (**D**) and (**E**), n=36 for both groups. Smears of total hemolymph collected from (**G**) *hml*>2x*GFP* and (**H**) *hml>HA-hipk^3M^+GFP* L3 larvae. (**I**) Quantification of mean number of hemocytes counted from genotypes in (**G**) and (**H**). Each data point represents the mean of 5 cell counts from one sample, *hml*>2x*GFP* (n=10 samples, n=50 cell counts) and *hml>HA-hipk^3M^+GFP* (n=10 samples, n=50 cell counts), P<0.0001. Scale bars equal 10μm.

We hypothesized that, similar to *hop^Tum-l^*, Hipk may increase the number of circulating hemocytes. We tested this by isolating the total hemolymph from third instar (L3) larvae (Fig. S1C), and determined that the mean number of hemocytes in each *hml>HA-hipk^3M^+GFP* sample was 197, compared to 39 per *hml*>2x*GFP* sample (Fig 1G-I). *hml>HA-hipk^3M^*+*GFP* samples often contained large aggregated clusters of hemocytes (Fig. S1B), a phenotype found in *hop^Tum-l^* hemocyte samples (Luo et al., 1997). These data suggest that the tumors induced by Hipk, like the ones induced by *hop^Tum-l^*, are derived from the hemocytes.

### hop^Tum-l^-induced lethality is rescued by reducing hipk

Extensive characterization of the *hop^Tum-l^* allele by others has shown that it can be utilized in lethality and tumor frequency assays to help in identifying novel JAK/STAT pathway components and regulators (Chen et al., 2014). We tested whether *hipk* could modify the *hop^Tum-l^* lethality phenotype (see Fig. 2F-H for ranking examples) (Rawlings et al., 2004; Yan and Luo, 1996). *hop^Tum-l^;;MKRS/TM6B* animals raised at 29^o^C were larval or early pupal lethal (Fig. 2A,B); 91% of pupae died in the early pupal stage, 9% died in the late pupal stage, and 0% of adults eclosed (Fig. 2E). Heterozygous reduction of *hipk (hop^Tum-1^;;hipk^4^/TM6B*) suppressed this phenotype (Fig. 2C,D); 23% of pupa died during early pupal development, and 76% died as pharate adults and 1% were able to eclose (Fig. 2E). Thus we infer that *hipk* is a positive regulator of the pathway since reducing *hipk* suppressed phenotypes caused by overactive JAK/STAT.

**Figure 2.**
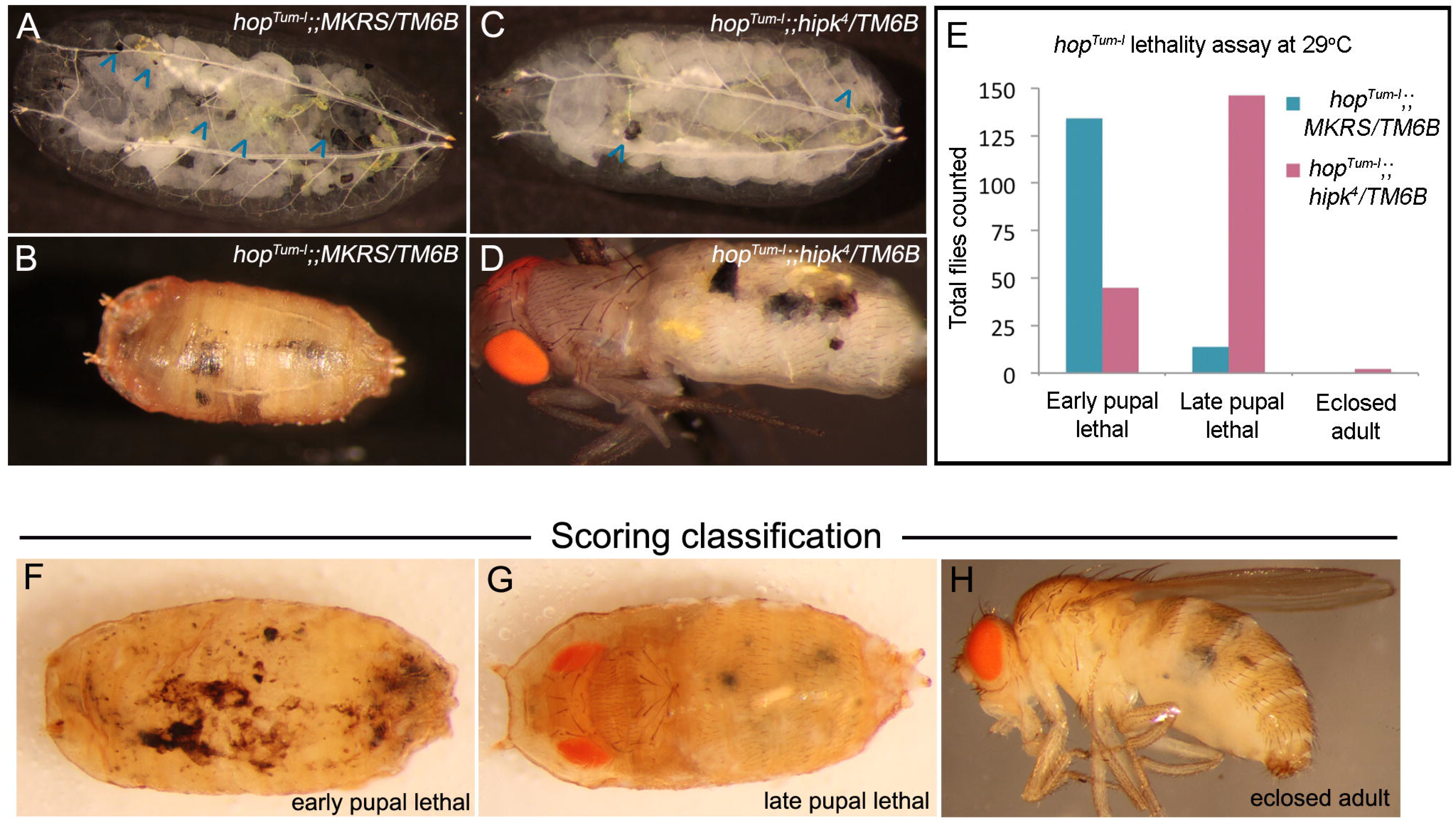
Heterozygous loss of *hipk* suppresses *hop^Tum-l^* induced lethality. (**A,B**)At 29°C *hop^Tum-l^* causes the formation of melanized tumors (**A**; arrowheads) and (**B**) results in larval and/or pupal lethality. (**C,D**) Heterozygous loss of *hipk* suppresses the tumor frequency (**C**; arrowheads) and (**D**) though some *hop^Tum-i‥^;;hipk*^4^/*TM6B* flies die in the early pupal stage, many reach the late pupal stage. (**E**) Quantification of the *hop^Tum-l^* lethality test in (**A-D**); *hop^Tum-l^;;MKRS/TM6B* (n=148) and *hop^Tum-l‥^;;hipk*^4^/*TM6B* (n=193). The *hop^Tum-l^* lethality assay was phenotypically ranked into three categories:(**F**) represents category ‘early pupal lethal’, where no adult structures are detectable,(**G**) represents the ‘late pupal lethal’ class, where adult structures are visible but the fly does not eclose, and (**H**) represents the class ‘eclosed adult’.

Since *hop^Tum-l^* tumors derive from hemocytes, we asked whether reduction of *hipk* within the hemocytes could rescue *hop^Tum-l^* lethality at 29°C. We expressed *UAS-hipk^RNAi^* with *hml-GAL4* in a *hop^Tum-l^* genetic background (*hop^Tum-l^/XorY;hml>hipk^RNAi^*), but did not observe a significant suppression (Fig. S2A-D). We reasoned that we were unable to obtain a rescue, possibly due to a combination of the strength of the *hop^Tum-l^* phenotype at 29°C, and weakness of *hml>UAS-hipk^RNAi^. hop^Tum-l^* is temperature sensitive, yielding a more severe phenotype at 29°C than at 25°C. We tested whether loss of *hipk* within hemocytes could rescue the *hop^Tum-l^* phenotype at 25°C. *hop^Tum-l^/Y;hml-GAL4/+* flies raised at 25°C exhibited a range of tumor frequencies: 15% of flies had more than 5 small to large tumors (class 1; Fig. S2E,H), 50% of flies had more than 5 small to medium tumors (class 2; Fig. S2F,H), and 35% of flies had less than 5 small tumors (class 3; Fig. S2G,H). Reducing *hipk*(*hop^Tum-l^/Y;hml-GAL4/UAS-hipk^RNAi^*) rescued the severity of *hop^Tum-l^* induced tumors; we observed 0% of flies in class 1, 29% of flies in class 2, and 71% of flies in class 3 (Fig. S2H). We conclude that *hipk* is required for the full severity of the *hop^Tum-l^* phenotype.

### *hipk* promotes JAK/STAT signaling downstream of *upd*

Cumulatively, our results suggest that *hipk* promotes JAK/STAT within the hemocytes. To further test *hipk’s* role with the JAK/STAT pathway we utilized the Stat92E transcriptional reporter 10xStat92E-GFP in L3 imaginal discs, which provides an accurate representation of endogenous pathway activity (Bach et al., 2007). Loss of *hipk* in somatic clones led to significant cell-autonomous reductions in 10xStat92E-GFP expression in wing and eye-antennal imaginal discs (Fig. 3B, Fig. S3B), while *dpp>HA-hipk^3M^* wing discs raised at 25°C, and at 29°C, caused elevated 10xStat92E-GFP (Fig. 3E, Fig. S3C). We found that expressing *UAS-HA-hipk^WT^-attP40* within *hipk^4^* clones could restore, and in some instances elevate 10xStat92E-GFP levels (Fig. 3C) indicating that the effect we see is due to Hipk expression. Next we tested if Hipk’s kinase activity was crucial for this effect on the JAK/STAT reporter. Expression of kinase dead Hipk (*UAS-HA-hipk^KD^-attP40*) within *hipk^4^* MARCM clones was unable to promote expression of 10xStat92E-GFP (Fig. 3D). Further support that Hipk is a positive regulator of JAK/STAT, we find that heterozygosity for *hipk* enhanced the small eye phenotype seen in *outstretched* (*os*; or Unpaired *upd*) mutants (Fig. S3F-J). Together, these results indicate that Hipk promotes JAK/STAT activity and is required for the proper output of the pathway in a cell autonomous fashion and kinase-dependent manner.

**Figure 3.**
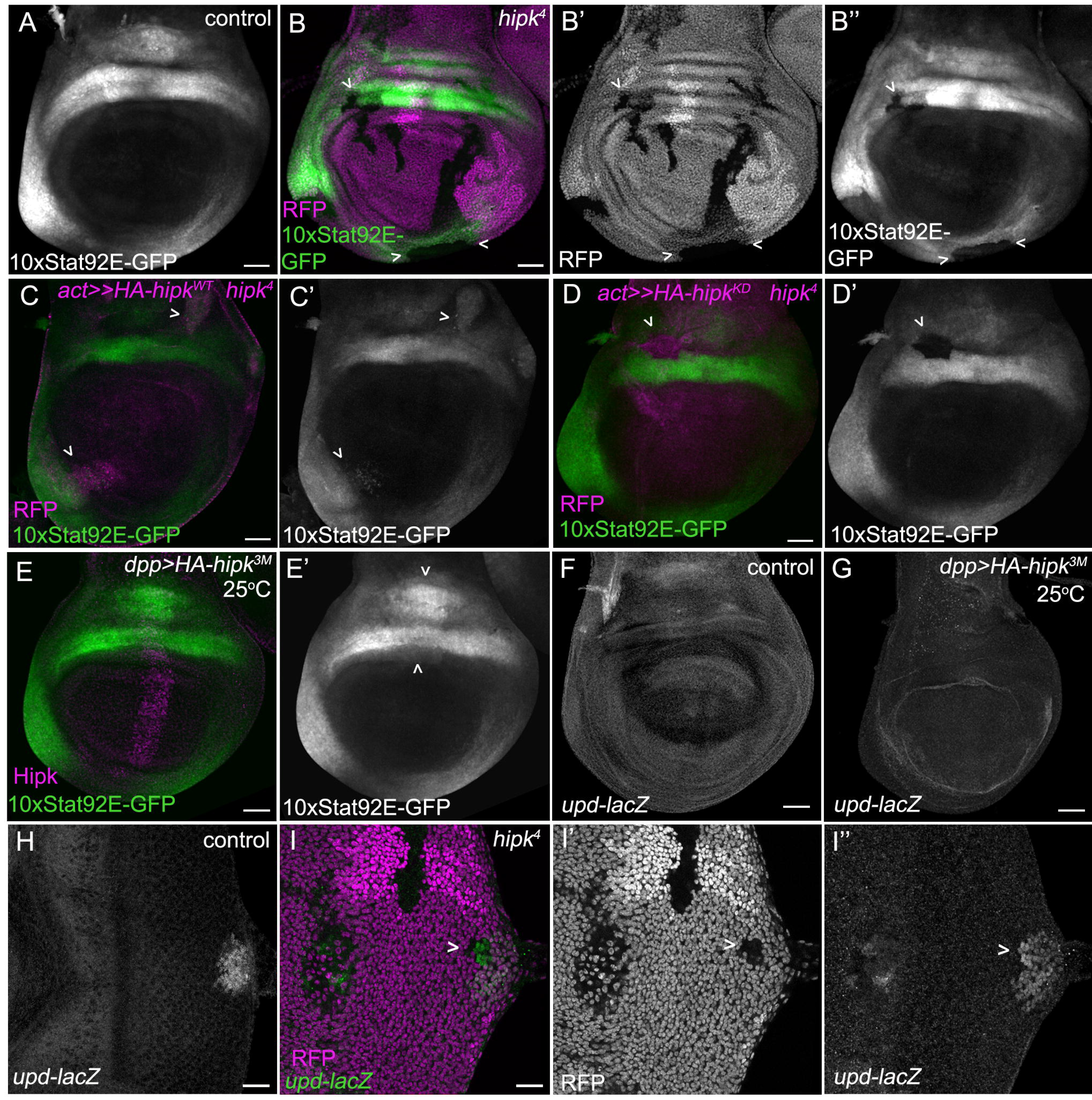
*hipk* promotes and is required for JAK/STAT signaling, downstream of *upd*. (**A**) A control L3 wing disc showing the expression domain of the reporter *10xStat92E-GFP*. (**B**) Expression of 10xStat92E-GFP is perturbed in *hipk*^4^ mutant clones marked by the absence of RFP (arrowheads) (n=20). (C) Expressing *UAS-HA-hipk^WT^* within *hipk*^4^ MARCM clones (*act≫HA-hipk^WT^*;*hipk*^4^) restores and can elevate 10xStat92E-GFP levels (n=10). (**D**) 10xStat92E-GFP levels are not restored within *act≫HA-hipk^KD^;hipk*^4^ clones (n=5). (**E**) Increases in 10xStat92E-GFP expression are observed in *dpp>HA-hipk^3M^* wing discs (arrowheads) raised at 25°C (n=20). (**F**) A control wing disc showing the expression domain of *upd-lacZ*. (**G**) At 25°C, *upd-lacZ* appears unchanged in *dpp>HA-hipk^3M^* wing discs (n=20). (**H**) *upd-lacZ* is expressed at the posterior center of the L3 eye-antennal control disc. (**I**) Loss of *hipk*, in negatively marked RFP clones, does not affect *upd-lacZ* (arrowhead) (n=20). Scale bars equal 10 μm.

To determine whether *hipk* promotes JAK/STAT activity upstream or downstream of Upd, we examined *upd-lacZ* expression in imaginal discs upon modulation of *hipk. upd-lacZ* is not prevalent in the L3 wing disc (Fig. 3F), but is expressed in cells at the posterior center of the L3 eye disc (Fig. 3H). Loss of *hipk* in the eye disc did not alter *upd* expression (Fig. 3I) and *upd* remained unchanged in *dpp>HA-hipk^3M^* wing discs raised at 25°C, conditions under which 10xStat92E-GFP is normally induced by Hipk (Fig. 3G, E'). However, *dpp>HA-hipk^3M^* wing discs raised at 29°C exhibited a small amount of *upd* up-regulation (Fig. S3D). While we observed that high levels of Hipk induce mild ectopic *upd* in the wing disc (Fig. S3D), past studies have shown that *hipk* promotes multiple signaling pathways, and this could represent an indirect up-regulation of *upd*. With this in mind, we conclude that *hipk* promotes JAK/STAT activity downstream of *upd*.

### Hipk physically interacts with Stat92E

Next we attempted to elucidate how Hipk was mediating its effects on the JAK/STAT pathway. Though partly cytoplasmic, Hipk primarily localizes to the nucleus (Kim et al., 1998). Because Stat92E is also found in the nucleus, we began testing for a physical Hipk-Stat92E interaction. We utilized a proximity ligation assay (PLA), which can detect whether two proteins of interest are less than 40nm apart *in vivo* (Söderberg et al., 2006). In *dpp>HA-hipk^1M^+MYC-Stat92E* wing discs we probed with HA and MYC antibodies and observed a positive PLA reaction (Figure 4C,D). While these are ectopically expressed proteins, it is clear from the mild disc phenotype that the expression levels are not extreme and that the interaction is unlikely to be due to protein saturation. Negative control discs (*dpp>HA-hipk^1M^+GFP*) that were probed against GFP and HA did not yield a PLA signal (Fig. 4A). These data suggest that Hipk and Stat92E come into close proximity in wing disc cells, though we cannot exclude the possibility that Hipk and Stat92E may come into close proximity as part of a protein complex.

**Figure 4.**
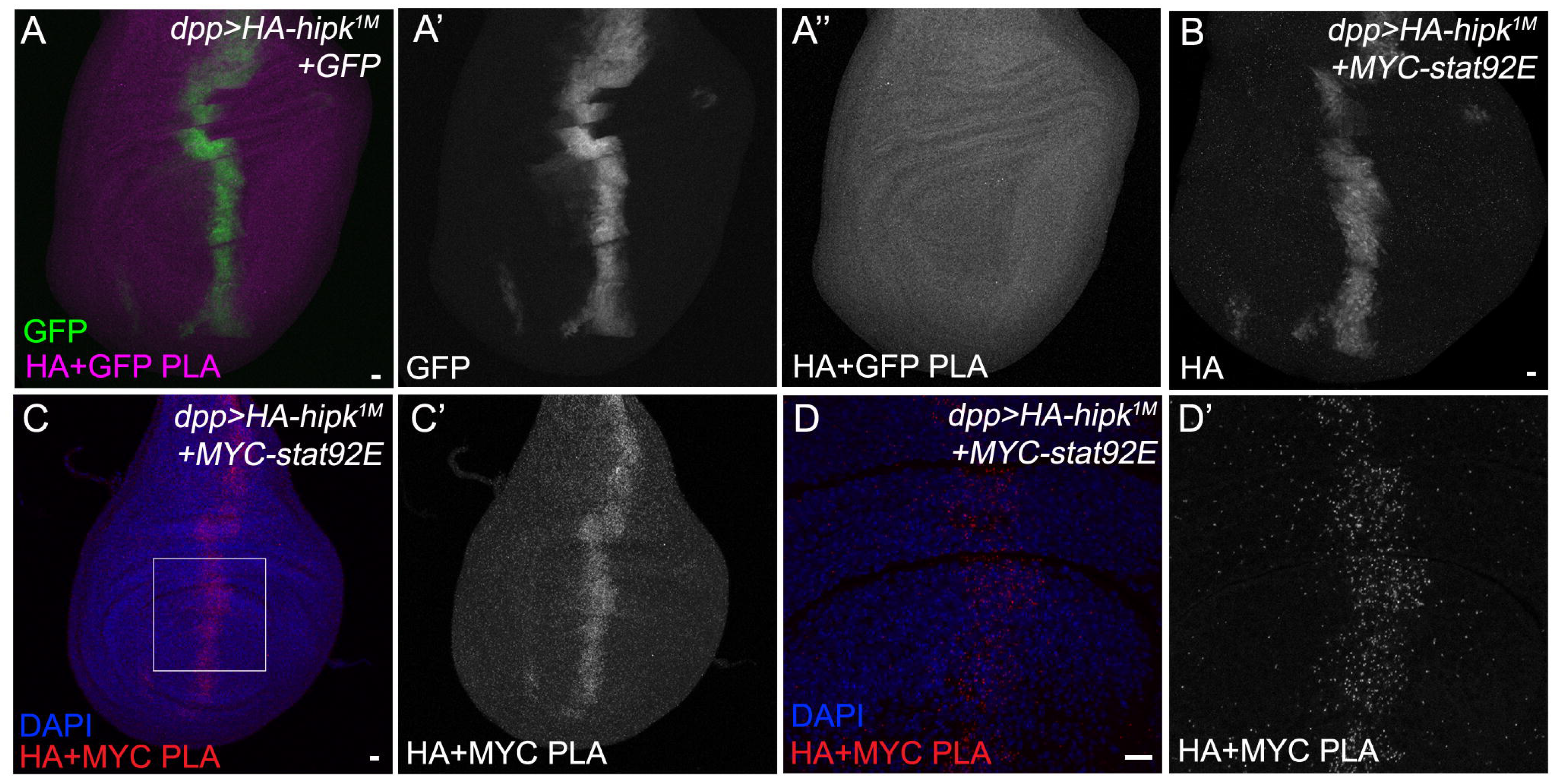
HA-Hipk and MYC-Stat92E physically interact in the wing imaginal disc. PLAs were performed on L3 wing imaginal discs. (**A**) A negative control disc probed with anti-HA and anti-GFP antibodies did not yield a positive signal (n=17). (**B**) *dpp>HA-hipk^1M^*+*MYC-stat92E* discs stained for HA show that Hipk is present in the same discs used for the PLA experiments. (**C,D**) A positive PLA signal is present along the *dpp* domain in *dpp>HA-hipk^1M^*+*MYC-stat92E* discs that were probed with HA and MYC (n=54). The boxed region in (**C**) represents the zoomed-in region in (**D**). Scale bars equal 10 m.

In summary, we present novel evidence that Hipk is an essential regulator of the JAK/STAT pathway in normal and tumorigenic processes and acts downstream of *upd* in a kinase-dependant manner. Output of JAK/STAT is perturbed upon loss of *hipk*, and increased Hipk induces hematopoietic tumors and elevated JAK/STAT activity. Further, we provide *in vivo* data that suggests a physical interaction between Hipk and Stat92E. Reports in vertebrate studies have found that an activated version of Hipk2 phosphorylates Stat3 (Matsuo et al., 2001; Ohtsu et al., 2007), and that Hipk2 is a potential drug target in treating Acute Myeloid Leukemia (Fleischmann et al., 2014). Future studies with help determine the precise mechanism of Hipk’s role in this pathway and could ultimately lead to new therapeutics used to treat human cancers.

## Methods

**Genetic crosses and fly stocks**: Flies were raised on standard media. Crosses were raised at 25°C unless otherwise noted. *10xstat92E-GFP* (BL#26197) (Bach et al., 2007), *UAS-eGFP* (BL#5431), *UAS-eGFP* (BL#5430), *UAS-P35* (BL#5072) (Hay et al., 1994), *hsflp^122^;;Ubi-RFP,FRT79* (made from BL#34498), *y^1^v^1^hop^Tum^/FM7c* (BL#8492; referred to as *hop^Tum-l^*), *act5c-GAL4/CyO* (BL#4414), *UAS-myr-RFP/CyO* (BL#7118) and *hml-GAL4* (BL#30139) were obtained from Bloomington Drosophila Stock Center, Bloomington, IN. *UAS-hipk^RNAi^* (VDRC#108254) was obtained from Vienna Drosophila Resource Center, Vienna, Austria. Also used were *dpp-GAL4/TM6B* (Staehling-Hampton et al., 1994), *os,y* (a gift from Norbert Perrimon), *w;UAS-Stat92E-Myc/Cyo,wg-lacZ* (a gift from Sol Sotillos) (Sotillos et al., 2013), *PD-lacZ* (a gift from Henry Sun; referred to as *upd-lacZ* hereon after) (Tsai and Sun, 2004), *ywhsflp,tub-GAL4, UAS-GFP, 6X MYC-NLS; UAS-y+;tub-GAL80,FRT2A/TM6B* (a gift from Gary Struhl), *ywhsflp^122^;sp/Cyo;TM2/TM6B, UAS-HA-hipk^1M^*, *UAS-HA-hipk^3M^*, *hipk*^4^, *FRT79/TM6B* (Lee et al., 2009a), *UAS-HA-hipk^K221R^-attP40* (kinase dead Hipk; hereafter referred to as *UAS-HA-hipk^KD^-attP40*) (Chen and Verheyen, 2012), and *UAS-HA-hipk^WT^-attP40* (made in this study). *act5c-GAL4/Cyo* and *UAS-myr-RFP/CyO* were recombined to generate *act5c-GAL4,UAS-myr-RFP/CyO*. *hipk*^4^, *FRT79/TM6B and10xstat92E-GFP/TM6B* were recombined to generate *hipk*^4^, *FRT79,10xstat92E-GFP/TM6B*.

**Generation of transgenic fly stocks**: DNA cloning was performed by Ziwei Ding of the SFU Molecular Biology Service Centre. pCMV-HA-Hipk (Lee et al., 2009a) was used as the source of HA-Hipk. The EcoRI site of pUASt-attB was mutated to a SmaI site, and HA-hipk^WT^ was inserted into this site. HA-hipk^WT^-attB was inserted into the attP40 locus generating the fly strain *UAS-HA-hipk^WT^-attP40* (Best Gene, Chino Hills, CA).

**Clonal analysis**: Somatic clones were generated by crossing *hsflp*^122^;;*Ubi-RFP,FRT79* to either *10XStat92E-GFP*;*hipk*^4^,*FRT79/TM6B*, or *upd-lacZ*;;*hipk*^4^,*FRT79/TM6B*. Progeny were heat shocked at 38 °C, 48 hours after egg laying for 90 minutes. MARCM clones were generated by crossing *ywhsflp*^122^;*act5c-GAL4,UAS-myr-RFP/CyO;tub-GAL80,FRT2A/TM6B* (RFP MARCM79) to either *hipk*^4^,*FRT79,10xstat92E-GFP/TM6B*, *UAS-HA-hipk^WT^-attP40;hipk*^4^,*FRT79,10xstat92E-GFP/SM6a~TM6B*, or *UAS-HA-hipk^KD^-attP40;hipk*^4^,*FRT79,10xstat92E-GFP/SM6a~TM6B*. Progeny were heat shocked at 38°C, 48 hours after egg laying for 90 minutes and were subsequently raised at 29°C.

**Immunocytochemistry and microscopy**: L3 imaginal discs were dissected and stained using standard protocols, and where possible we analyzed equal to or greater than 20 discs per genotype; the exception to this was the MARCM experiments in Fig. 3 and S3 which were particularly sickly and thus had lower n-values. Detailed information regarding antibodies used and methods of microscopy can be found in the Supplementary Materials.

**Eye size comparison for *os* assay**: 10 images were acquired for *TM3/TM6B, hipk*^4^/*TM6B, os;;MKRS/TM6B*, and *os;;hipk*^4^/*TM6B* adult eyes. The area of each eye was measured in pixels using Photoshop, and the values were subjected to a student’s t-test.

***hop^Tum-l^* lethality and tumor frequency assays**: The lethality assay in Fig. 2 was performed by crossing 50 females and 15 males from each stock (*hop^Tum-l^;;MKRS/TM6B* and *hop^Tum-l^;;hipk*^4^/*TM6B*) in a bottle and raising flies at 29°C. After 11 days, all pupae were removed from the walls of the bottles and were ranked as either ‘early pupal lethal’ (had no recognizable adult structures), ‘late pupal lethal’ (pharate adults), or ‘eclosed adult’ (see examples of each rank in Fig. 2F-H). The lethality assay in Fig. S2A-D was performed by crossing 8 females (*hop^Tum-l^/(FM7);hml-GAL4*) to 6 males (either *w^1118^*/*Y* or *UAS-hipk^RNAi^*) in a vial and raising flies at 29°C. Progeny were scored using the same methods as the previous lethality assay (for scoring examples see Fig. S2A-C). The tumor frequency assay in Fig. S2E-H was performed by crossing 8 females (*hop^Tum-l^/(FM7);hml-GAL4*) to 6 males (either *w^1118^*/*Y* or *X/Y;UAS-hipk^RNAi^*) in a vial and raising flies at 25°C. After 13 days, male progeny were scored into the following classifications: ‘class 1’ (flies had greater than 5 tumors ranging in size from small to large), ‘class 2’ (more than 5 small to medium tumors were present), and ‘class 3’ (less than 5 small tumors were present) (see examples of each class in Fig. S2E-G).

**Hemocyte counts**: Prior to hemolymph collection, L3 larvae were washed thoroughly with 1X PBS, dried, and placed in a glass dissection well containing 5 μL of 1X PBS. The larval cuticle was carefully punctured with forceps and hemolymph was allowed to drain into the well. Each sample contains the hemolymph from two larvae in 5uL of 1X PBS. Hemolymph was then smeared onto a poly-D-lysine coated slide and air-dried. Cell smears were washed with 3.7% formaldehyde for 5 minutes, washed with PBS, and stained with DAPI. For each sample (n=10), 5 cell counts were performed and means of the 5 cell counts were plotted; values were subjected to a student’s t-test.

**Proximity Ligation Assay (PLA)**: PLA was performed on L3 wing discs according to standard protocols (Wang et al., 2014) with the following exceptions: discs were fixed with 4% formaldehyde for 15 minutes and discs were blocked with 1% normal donkey serum. A PLA against HA and GFP was used as a negative control. A subset of the discs were stained for HA to ensure that Hipk was expressed.

## Acknowledgements

We are grateful to N. Perrimon, H. Sun, S. Sotillos, Bloomington Drosophila Stock Center (NIH P400D018537), and Developmental Studies Hybridoma Bank for providing fly strains and antibodies. Also, we thank Z. Ding for help in creating the HA-hipkWT-attP40 construct, and A. Kadhim for help with crosses. This work was funded by an operating grant from the Canadian Institutes of Health Research. NBW was supported by a PGS-M fellowship award from N.S.E.R.C.

## Author contributions

Conception and design JB, NW, EV. Acquisition of data JB, NW. Analysis and interpretation of data JB, NW, EV. Drafting or revising the article JB, NW, EV.

